# Structure Reveals Homology in Elevator Transporters

**DOI:** 10.1101/2023.06.14.544989

**Authors:** Noah Trebesch, Emad Tajkhorshid

## Abstract

The elevator transport mechanism is one of the handful of canonical mechanisms by which transporters shuttle their substrates across the semi-permeable membranes that surround cells and organelles. Studies of molecular function are naturally guided by evolutionary context, but until now this context has been limited for elevator transporters because established evolutionary classification methods have organized them into several apparently unrelated families. Through comprehensive examination of the pertinent structures available in the Protein Data Bank, we show that 62 elevator transporters from 18 families share a conserved architecture in their transport domains consisting of 10 helices connected in 8 topologies. Through quantitative analysis of the structural similarity, structural complexity, and topologically-corrected sequence similarity among the transport domains, we provide compelling evidence that these elevator transporters are all homologous. Using our analysis, we have constructed a phylogenetic tree to enable quantification and visualization of the evolutionary relationships among elevator transporters and their families. We also report several examples of functional features that are shared by elevator transporters from different families. Our findings shed new light on the elevator transport mechanism and allow us to understand it in a far deeper and more nuanced manner.

## 1 Introduction

Transport of substances across the mostly impermeable membranes that surround cells and cellular organelles is a fundamental biological process, and transporters are one of the major classes of proteins that mediate this process. Because of their central biological role as cellular gatekeepers, transporter malfunction is often associated with disease, and transporters constitute a major target for pharmacological intervention.^1^ The function of transporters is underpinned by their structural dynamics, and detailed understanding of these structural dynamics hold deep implications for both biomedicine and basic science.

During their functional cycles, transporters achieve substrate transport by structurally transitioning between their inward facing (IF) and outward facing (OF) conformational states. This process changes the accessibility of the substrate binding site from one side of the membrane to the other and is known as the alternating access mechanism.^2^ The structural details by which the alternating access mechanism is achieved are complex and vary widely from one transporter to another, but a handful of canonical transport mechanisms have emerged as more structures have been solved,^3^ one of which is referred to as the “elevator” transport mechanism.

Elevator transporters are composed of one or more functional units called protomers that each consist of a transport domain and a scaffold domain (sometimes also referred to as core and gate domains, respectively). In the elevator transport mechanism, the accessibility of the substrate binding site is alternated from one side of the membrane to the other through characteristic translational and rotational motions of the transport domain relative to the scaffold domain (Fig. 1).^4^ As more structures of transporters have been solved, they have revealed that several families of transporters operate via the elevator mechanism.^5^

**Figure 1:**
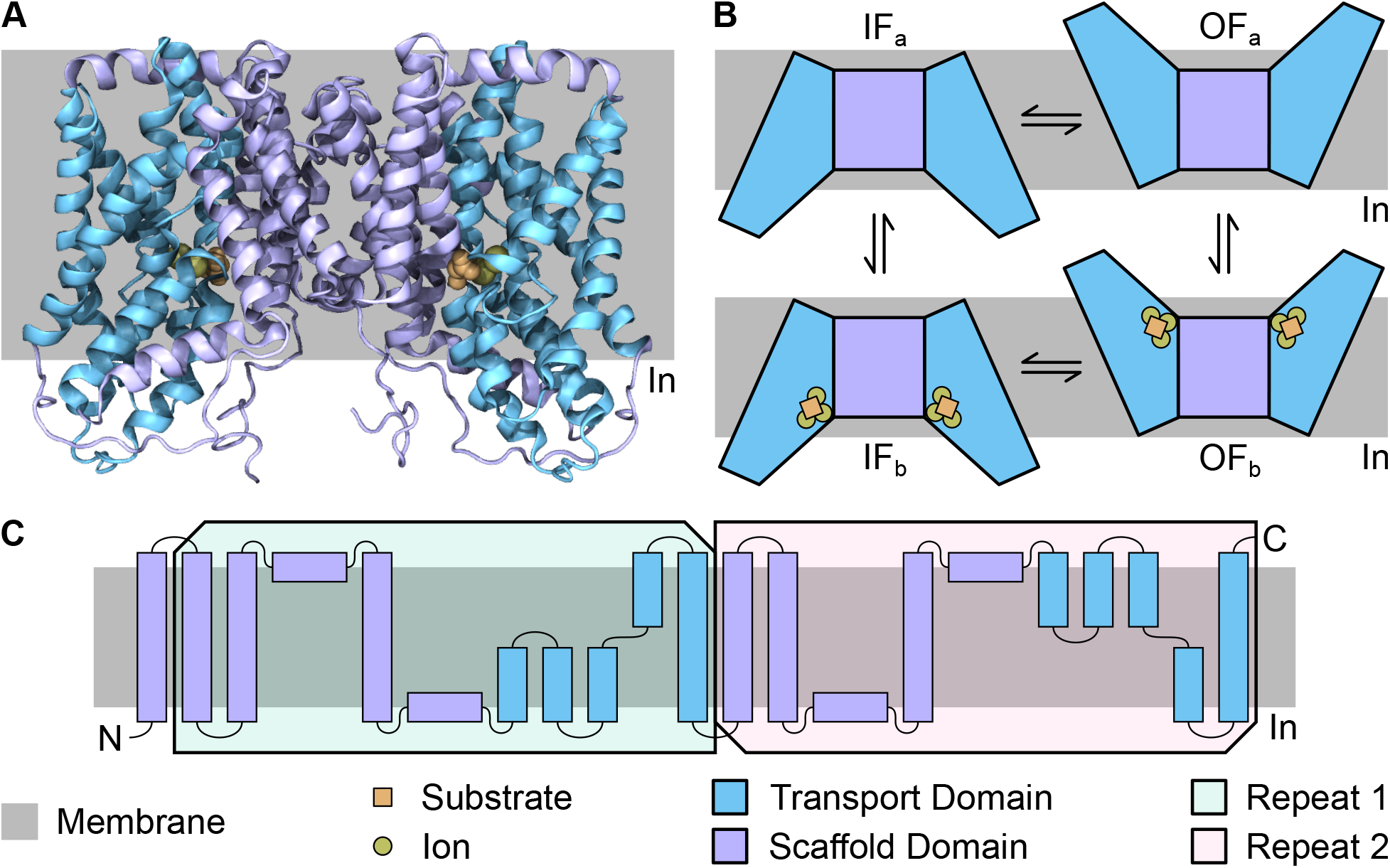
Structural and mechanistic features of VcINDY, a representative elevator transporter. **(A)** Atomistic structure of VcINDY (PDB `5UL7`) in an IF state bound to substrate (succinate) and ions (Na^+^). VcINDY is a homodimer, and as in all other multimeric elevator transporters, the scaffold domain is also responsible for oligomerization. **(B)** The transport cycle of VcINDY. The transporter starts with a transition from its apo IF (IF_a_) state to its apo OF (OF_a_) state, which involves large-scale rigid-body rotation and translation (i.e., elevator-like motion) of the transport domains relative to the scaffold domain. Once in the OF_a_ state, substrate and ions bind to VcINDY, generating a bound OF (OF_b_) state. The transport domains then undergo large-scale elevator-like motions again, generating a bound IF (IF_b_) state. Finally, substrate and ions unbind, returning VcINDY to its IF_a_ state. For simplicity, both protomers of VcINDY are shown in the same state in this figure, but they are thought to operate independently. **(C)** Topology of a VcINDY protomer. As with most elevator transporters, the topology of VcINDY contains so-called “inverted repeats,” highlighted by the pink and green boxes in the figure. For each repeat, the topology is identical, except the placement and N-to-C orientation of the helices relative to the membrane are inverted.

Evolution offers a natural framework through which investigation and understanding of protein function can be viewed. Functional details from one protein are more likely to be shared by closely related proteins than unrelated or more distantly related ones. Indeed, the first characterization of a family of proteins is usually viewed as inherently more significant than corroborative characterization of additional proteins within a well studied family. Accordingly, the establishment of meaningful protein families is of critical importance in the investigation and understanding of protein function.

The classification system most widely used to group homologous transporters into families and superfamilies is the Transporter Classification Database (TCDB).^6^ TCDB uses a sophisticated sequence-based approach to establish statistically significant evidence of homology among transporters. Another widely used classification system for transporters, as well as for other proteins, is the Pfam system,^7^ which uses a hidden Markov model (HMM)-based approach to establish protein homology and which has largely reproduced TCDB’s original transporter families. The TCDB and Pfam approaches of establishing homology are powerful but also have their own fundamental limitations.

During protein evolution, it is known that protein architecture is conserved far longer than amino acid sequence and even protein topology, and conservation of architecture is considered the best and only evidence capable of establishing distant evolutionary relationships.^8–10^ Here, we use the term “architecture” to refer to the three dimensional arrangement of a protein’s secondary structural elements relative to one another, and we use the term “topology” to refer to the connectivity of those secondary structural elements within the protein’s polypeptide chain as well as their N-to-C orientations with respect to one another. Standard sequence alignment algorithms are not capable of producing meaningful global alignments when the topologies of related proteins have diverged, so the sequence-based approaches of TCDB and Pfam are not capable of establishing homology among these more distantly related proteins.

Pioneering work has previously noted striking structural similarity among elevator transporters.^3, 11^ We also noted this similarity and have sought to characterize it quantitatively and comprehensively, focusing here primarily on the transport domains of elevator transporters. After comprehensive examination of the elevator transporter structures that were available in the Protein Data Bank (PDB)^12^ as of May 10, 2023, we determined that 62 of them from 18 TCDB families share a conserved transport domain architecture. Through detailed analysis of the architectures, topologies, and sequences of these transport domains, we found compelling quantitative evidence that these elevator transporters are homologous to each other. From these results, we generated a phylogenetic tree of the elevator transporters, and to illustrate the tree’s practical utility, we identified several preliminary examples of fundamental functional features shared by transporters from adjacent clades in the tree.

## 2 Results

### 2.1 Structurally Characterized Elevator Transporters

Through comprehensive visual examination of the elevator transporter structures deposited in the PDB, we identified 62 elevator transporters that share a conserved architecture within their transport domains (Table 1). Besides the conserved architecture and mechanism, there is significant diversity in this collection of transporters. The transporters belong to 18 TCDB families, and 12 of these families belong to 5 TCDB superfamilies. The transporters are from a wide variety of organisms; 4 are are from archaea, 23 are from bacteria, and 35 are from eukarya (17 of which are from humans). They operate in a variety of oligomeric states; 7 operate as monomers, 43 as dimers, and 12 as trimers. Finally, they transport a wide variety of substrates along with various ions in different stoichiometries, and some have even lost their transport function (specifically hPrestin, MuPrestin, and TtPrestin).

**Table 1:** Elevator transporters included in this study. The PDB column follows the format “PDB Identifier - Chain.” The TCDB Classification column follows the format “Super-family : Family.” The Topology column refers specifically to the topology of the transport domain (see Fig. 3).

| **Protein** | **PDB** | **Organism** | **TCDB**<br>**Classification** | **Oligomeric**<br>**State** | **Topology** |
| --- | --- | --- | --- | --- | --- |
| Glt<sub>Ph</sub> | 7RCP-A | Archaeon | : DAACS | Trimer | Glt<sub>Ph</sub> |
| Glt<sub>Tk</sub> | 5DWY-A | Archaeon | : DAACS | Trimer | Glt<sub>Ph</sub> |
| EAAT1 | 5LM4-A | Human | : DAACS | Trimer | Glt<sub>Ph</sub> |
| hEAAT2 | 7XR4-A | Human | : DAACS | Trimer | Glt<sub>Ph</sub> |
| hEAAT3 | 8CUA-A | Human | : DAACS | Trimer | Glt<sub>Ph</sub> |
| ASCT1 | 7P4I-A | Human | : DAACS | Trimer | Glt<sub>Ph</sub> |
| ASCT2 | 7BCT-A | Human | : DAACS | Trimer | Glt<sub>Ph</sub> |
| VcINDY | 5UL9-A | Bacterium | IT : DASS | Dimer | vcCNT |
| LaINDY | 6WTW-A | Bacterium | IT : DASS | Dimer | vcCNT |
| NaCT | 7JSK-A | Human | IT : DASS | Dimer | vcCNT |
| MtrF | 4R1I-A | Bacterium | IT : AbgT | Dimer | vcCNT |
| YdaH | 4R0C-A | Bacterium | IT : AbgT | Dimer | vcCNT |
| HiSiaQM | 7QE5-A | Bacterium | IT : TRAP-T | Monomer | vcCNT |
| PpSiaQM | 7QHA-B | Bacterium | IT : TRAP-T | Monomer | vcCNT |
| HvYS1 | 7WST-A | Plant | : OPT | Dimer | HvYS1 |
| bcChbC | 3QNQ-A | Bacterium | PTS-GFL : Glc | Dimer | bcChbC |
| bcMalT | 5IWS-A | Bacterium | PTS-GFL : Glc | Dimer | bcChbC |
| vcCNT | 4PD6-A | Bacterium | : CNT | Trimer | vcCNT |
| CNT<sub>NW</sub> | 5L26-A | Bacterium | : CNT | Trimer | vcCNT |
| hCNT3 | 6KSW-A | Human | : CNT | Trimer | vcCNT |
| StOAD | 6IWW-A | Bacterium | CPA : NaT-DC | Trimer | KpCitS |
| KpCitS | 5XAS-A | Bacterium | : 2-HCT | Dimer | KpCitS |
| SeCitS | 5A1S-A | Bacterium | : 2-HCT | Dimer | KpCitS |
| SbtA | 7EGK-A | Bacterium | : SBT | Trimer | SbtA |
| PIN1 | 7Y9T-A | Plant | BART : AEC | Dimer | SbtA |
| AtPIN3 | 7WKW-A | Plant | BART : AEC | Dimer | SbtA |
| PIN8 | 7QP9-A | Plant | BART : AEC | Dimer | SbtA |
| ASBT<sub>NM</sub> | 3ZUX-A | Bacterium | BART : BASS | Monomer | EcNhaA |
| ASBT<sub>Yf</sub> | 6LGV-A | Bacterium | BART : BASS | Monomer | EcNhaA |
| RnNTCP | 7VAF-A | Rat | BART : BASS | Monomer | EcNhaA |
| BtNTCP | 7VAE-A | Cow | BART : BASS | Monomer | EcNhaA |
| NTCP | 7ZYI-A | Human | BART : BASS | Monomer | EcNhaA |
| EcNhaA | 7S24-A | Bacterium | : NhaA | Dimer | EcNhaA |
| StNhaA | 7A0W-A | Bacterium | : NhaA | Dimer | EcNhaA |
| TtNapA | 5BZ3-A | Bacterium | CPA : CPA2 | Dimer | EcNhaA |
| MjNhaP1 | 4CZB-A | Archaeon | CPA : CPA1 | Dimer | EcNhaA |
| PaNhaP | 4CZ8-A | Archaeon | CPA : CPA1 | Dimer | EcNhaA |
| NHE1 | 7DSW-A | Human | CPA : CPA1 | Dimer | EcNhaA |
| NHE3 | 7X2U-A | Human | CPA : CPA1 | Dimer | EcNhaA |
| NHE9 | 6Z3Z-A | Horse | CPA : CPA1 | Dimer | EcNhaA |
| BbNHA2 | 7P1J-A | Bison | CPA : CPA1 | Dimer | EcNhaA |
| NHA2 | 7B4L-A | Human | CPA : CPA1 | Dimer | EcNhaA |

| Protein | PDB | Organism | TCDB<br>Classification | Oligomeric<br>State | Topology |
| --- | --- | --- | --- | --- | --- |
| UraA | 5XLS-A | Bacterium | APC : NCS2 | Dimer | UraA |
| PurT<sub>Cp</sub> | 7TAK-A | Bacterium | APC : NCS2 | Dimer | UraA |
| UapA | 5I6C-A | Fungus | APC : NCS2 | Dimer | UraA |
| MmSVCT1 | 7YTW-A | Mouse | APC : NCS2 | Dimer | UraA |
| Bor1 | 5L25-A | Plant | APC : AE | Dimer | UraA |
| NDCBE | 7RTM-A | Rat | APC : AE | Dimer | UraA |
| bAE1 | 8D9N-A | Cow | APC : AE | Dimer | UraA |
| AE1 | 7UZ3-C | Human | APC : AE | Dimer | UraA |
| hAE2 | 8GVC-A | Human | APC : AE | Dimer | UraA |
| NBCe1 | 6CAA-A | Human | APC : AE | Dimer | UraA |
| BicA | 6KI1-A | Bacterium | APC : SulP | Dimer | UraA |
| AtSULTR4;1 | 7LHV-A | Plant | APC : SulP | Dimer | UraA |
| SLC26Dg | 5DA0-A | Bacterium | APC : SulP | Dimer | UraA |
| SLC26A2 | 7XLM-A | Human | APC : SulP | Dimer | UraA |
| MmSLC26A9 | 6RTC-A | Mouse | APC : SulP | Dimer | UraA |
| SLC26A9 | 7CH1-A | Human | APC : SulP | Dimer | UraA |
| MuPrestin | 7SUN-A | Gerbil | APC : SulP | Dimer | UraA |
| TtPrestin | 7S8X-A | Dolphin | APC : SulP | Dimer | UraA |
| hPrestin | 7LGU-A | Human | APC : SulP | Dimer | UraA |
| Pendrin | 7WK1-A | Mouse | APC : SulP | Dimer | UraA |

### 2.2 Conserved Architecture of Elevator Transport Domains

Within the transport domains of elevator transporters, we identified a conserved architecture consisting of 10 helices (Fig. 2). To simplify our discussion, we refer to the side of the transport domain that interacts with the scaffold domain as the “front” of the domain and the opposite side (which interacts with the membrane) as the “back” of the domain. We further refer to the half of the domain embedded in the outer leaflet of the membrane as the “upper” half, and we refer to the half of the domain embedded in the inner leaflet as the “lower” half.

**Figure 2:**
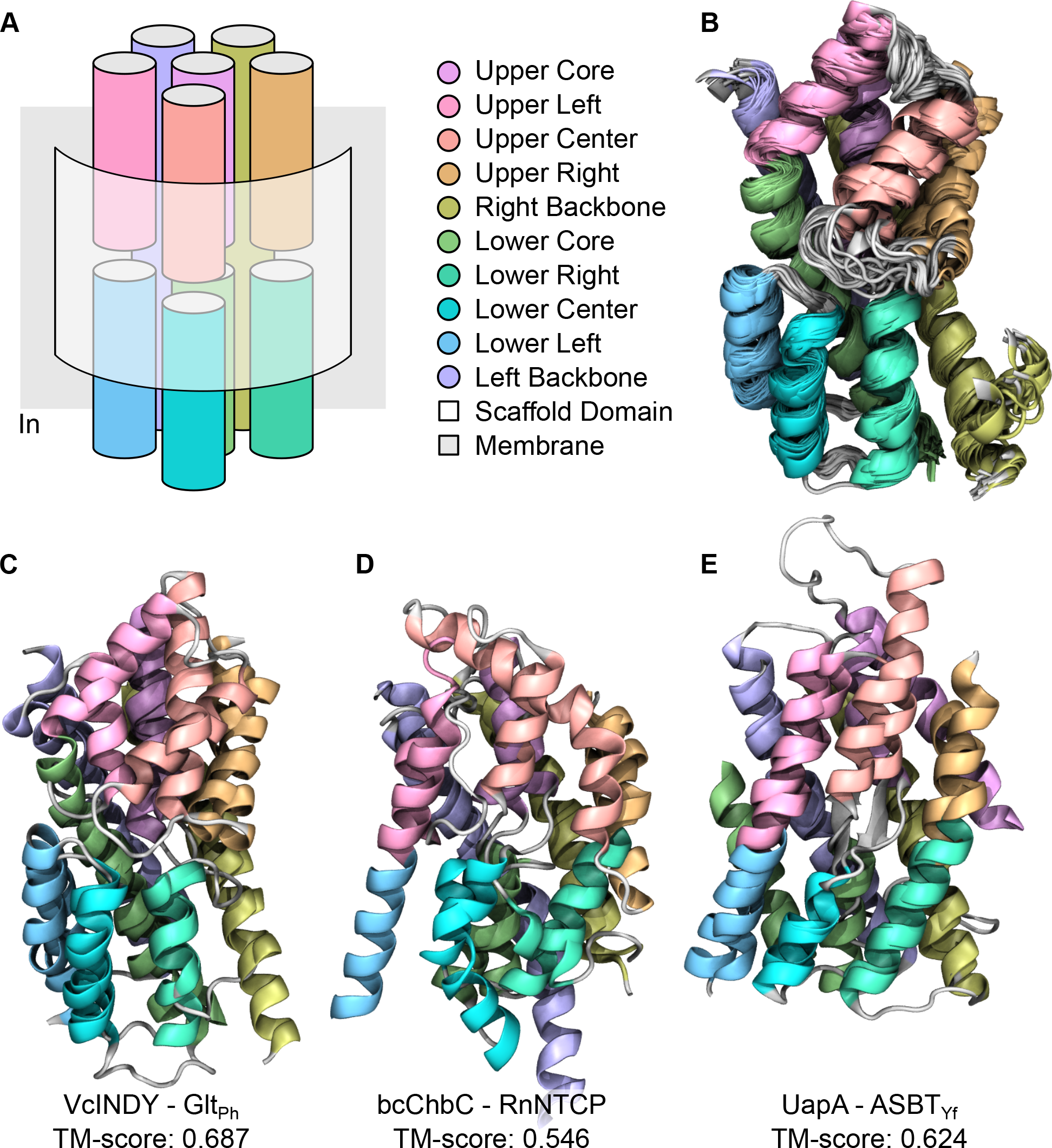
The conserved architecture of the transport domains of elevator transporters. **(A)** Simplified schematic of the conserved architecture, with unique names and colors assigned to each of the 10 constituent helices. We refer to the side of the transport domain facing the scaffold domain as the “front” side, and we refer to the half of the transport domain embedded in the outer leaflet of the membrane as the “upper” half. **(B)** Superposition of the transport domains from all Glt_Ph_ structures in the PDB (122 protomers from 44 PDBs). Of all elevator transporters, Glt_Ph_ is by far the most comprehensively structurally characterized, and it has been captured in a wide variety of functional states. This superposition illustrates the rigidity of the transport domain within the elevator mechanism, which we rely on to make meaningful structural comparisons between different transport domains. **(C)** Superposition of the transport domains of Glt_Ph_ (PDB `7RCP-A`) and VcINDY (PDB `5UL9-A`), a representative structural comparison. The transport domain helices of Glt_Ph_ and VcINDY are generally the same length and lie on top of one another, though there is some divergence in the positions of the upper left (pink) and upper center (red) helices. **(D)** Superposition of the transport domains of bcChbC (PDB `3QNQ-A`) and Rn-NTCP (PDB `7VAF-A`), which illustrates a notable example of variation from the conserved architecture. Specifically, bcChbC is missing its lower left (blue) helix, a feature common to all members of the PTS-GFL : Glc family. **(E)** Superposition of the transport domains of UapA (PDB `5I6C-A`) and ASBT_Yf_ (PDB `6LGV-A`), which illustrates another notable example of architectural variation. In UapA, the upper core (mauve) and lower core (green) helices have actually fallen out of the core, though they still overlap with the corresponding helices of ASBT_Yf_. This alternative arrangement of the core helices is common to all elevator transporters of the APC superfamily.

Using this terminology, the conserved architecture of the transport domains consists of two long helices (the “left backbone” and “right backbone” helices) at the back of the domain which always extend from the top of the membrane to the bottom. There are three upper (“upper left,” “upper center,” and “upper right”) and three lower (“lower left,” “lower center,” and “lower right”) helices that extend from one side of the membrane to its midplane and which interact with the scaffold domain. Finally, there are two more helices (the “upper core” and “lower core” helices) which also extend only halfway through the membrane and which are usually packed between the helices at the front and back of the domain. With this architectural annotation defined, we manually generated a list of the residues that make up each of the conserved transport domain helices from all elevator transporters for use in further analysis (Table S1).

Within the elevator transport domains, variations in this conserved architecture were generally small, and the conserved helices generally lay on top of one another when the domains were superposed (Fig. 2C), regardless of the transporter’s functional state (IF, OF, or occluded). However, there were a couple of notable exceptions. Specifically, transporters in the PTS-GFL : Glc family were missing their lower left helices (Fig. 2D). Additionally, the core helices of the elevator transporters in the APC superfamily “fell out” of the core of the domain, though parts of the APC core helices still overlapped with the core helices from other elevator transporters (Fig. 2E).

### 2.3 Topology of Elevator Transport Domains

Within the transport domains of elevator transporters, we observed 8 distinct topologies, which we named after the first published protein structure that exhibited each topology (Fig. 3). So-called “inverted repeats”^13^ (i.e., parts of a protein which have identical topologies, except the N-to-C orientations of the topologies are inverted with respect to one another, Fig. 1C) are observable in six of these topologies, while they have been lost in the Glt_Ph_ and bcChbC topologies. Notably, the SbtA and EcNhaA topologies are identical to each other, except the N-to-C order of their repeats are swapped relative to each other.

**Figure 3:**
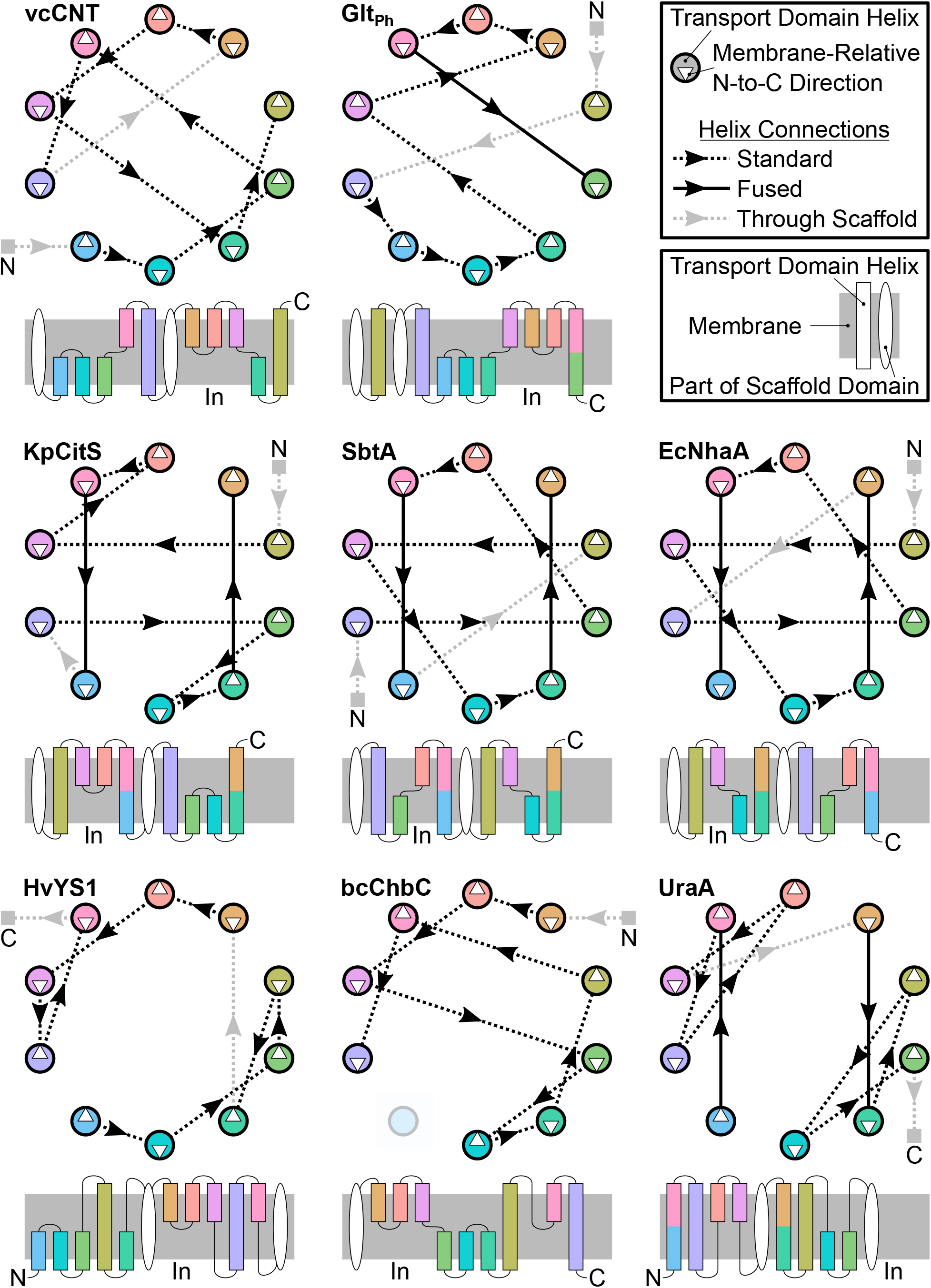
The varied topologies of the transport domains of elevator transporters. We found that the transport domains of elevator transporters adopt 8 distinct topologies, and we named each topology after the first published transporter structure that exhibited that topology. For each topology, the bottom schematic shows a traditional topology diagram, while the top schematic shows an alternative representation that better highlights similarities in helix connectivity with respect to the conserved architecture. Helices are colored according to the architectural annotation presented in Fig. 2.

In several topologies, two helices are fused together. Specifically, in the KpCitS, SbtA, EcNhaA, and UraA topologies, the upper left (pink) helix is fused to the lower right (blue) helix, and the upper right (orange) helix is fused to the lower right (turquoise) helix. In the Glt_Ph_ topology, the upper left (pink) helix is fused to the lower core (green) helix. Where two helices are fused, we note that there is often a kink, which we used to help determine where to split the helix for our architectural annotation.

### 2.4 Evidence of Homology

In proteins that appear structurally similar, distinguishing between homology produced by divergent evolution and analogy produced by convergent evolution is challenging, especially for membrane-embedded proteins which have a limited fold space.^14, 15^ Generally, proteins that are structurally complex and share rather similar structure, sequence, and function are more likely to be homologous than proteins that are rather simple and are less similar in their structure, sequence, and function.^10, 16^ Here, we put these qualitative guidelines in a quantitative framework and use it to suggest that the elevator transporters analyzed here are homologous.

As the primary basis for our analysis, we performed comprehensive pairwise structural and sequence comparisons of the elevator transport domains using TM-align.^17^ To generate meaningful global comparisons of the transport domains with TM-align, we used as input modified PDB files that corrected for topological differences between the transport domains (see Methods). On Zenodo (doi.org/10.5281/zenodo.8040643), we have made available a comprehensive set of PDB files that capture all of the pairwise structural superpositions produced by TM-align, along with the topology-corrected PDB files input to TM-align and the log files it produced. Heat maps of the resulting pairwise TM-scores^18^ and sequence identities are shown in Fig. 4, and we discuss them below in more detail.

**Figure 4:**
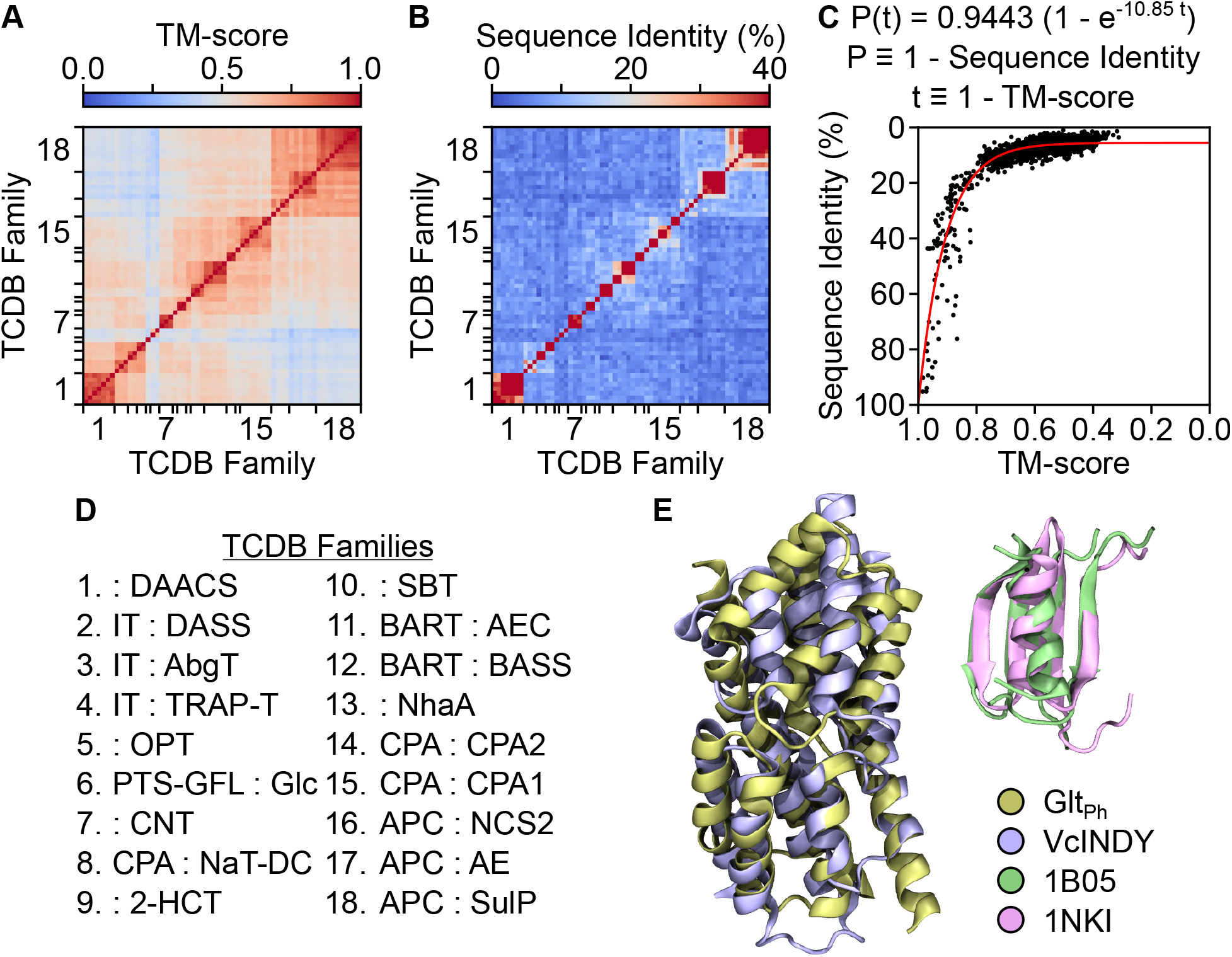
Evidence of homology among elevator transporters. **(A)** Heat map of the TM-scores resulting from the pairwise structural superpositions of elevator transport domains. In the plot, elevator transporters are grouped together by TCDB family, and tick marks indicate the start and end of each family. The mapping between the number assigned to each TCDB family and its name is found in (D). Past analysis has indicated that homologous proteins generally have a TM-score greater than or equal to 0.5, and all TCDB families are connected (directly or indirectly) by pairwise TM-scores that exceed this threshold. **(B)** Heat map of the sequence identities derived from the pairwise structural superpositions of elevator transport domains. A sequence identity of 20% is sometimes used to indicate homology between transporters, and this threshold is generally not met, even between transporters in the same TCDB family whose homology is well established. **(C)** Plot demonstrating the relationship between sequence identity and structural similarity in elevator transport domains. We obtained a nonlinear least-squares fit to our data using SciPy^23^ (specifically, we used the `optimize.least_squares` function with a `soft_l1` loss function and an `f_scale` of 0.1). The functional form of the fit is derived from a simple model of sequence divergence during evolution, suggesting that the sequence identity present in the transport domains is consistent with the idea that the elevator transporters are homologous. **(D)** Mapping of the numbers representing TCDB families in (A) and (B) to the TCDB family names. **(E)** Comparison of the structural complexity of a representative transport domain pair (Glt_Ph_, PDB `7RCP-A`, and VcINDY, PDB `5UL9-A`) with the structural complexity of a representative protein analog pair from the MALISAM database. Elevator transport domains are composed of 10 helices arranged in a complex architecture, while the protein motifs in the MALISAM database are composed of only a few *α*-helices and/or *β*-strands arranged in simple architectures.

#### 2.4.1 Structural Similarity

The Structural Classification of Proteins (SCOP)^19, 20^ and Class, Architecture, Topology, Homology (CATH)^21^ databases seek to hierarchically classify proteins by their structural and evolutionary relationships. At the highest level, both databases group together proteins with similar principal secondary structural content, and at the lowest level, they group together proteins that they have judged to be homologous through a combination of automated processes and extensive manual curation. Coverage by these databases of structures in the PDB is extensive but not comprehensive, and most elevator transporters have not yet been included.

Nevertheless, the SCOP and CATH databases offer valuable insight into the level of structural similarity present in homologous proteins, and the developers of TM-score and TM-align have analyzed these databases to assign meaning to different values of TM-score.^22^ Specifically, their structural analysis of protein families indicates that pairs of homologous proteins generally have a TM-score of 0.5 or higher, while random pairs of unrelated proteins generally have a TM-score of 0.3 or lower. Pairwise TM-scores of 0.5 or higher link together all of the elevator transport domains we have analyzed (Fig. 4A), suggesting that they are structurally similar enough to indicate homology.

#### 2.4.2 Sequence Similarity

Solute carrier (SLC) genes (which encode all human elevator transporters) are grouped into homologous families by the Human Genome Organization (HUGO) Gene Nomenclature Committee (HGNC) if their members share 20% or greater sequence identity.^1^ Pairwise sequence identity among elevator transport domains is quite low and generally below the simple HGNC SLC cutoff for homology (Fig. 4B). However, this result does not necessarily suggest that the transporters are not homologous as sequence identity is low even between members of TCDB families and superfamilies, whose homology has already been well established. A more sophisticated approach is required to determine whether homology is supported at the sequence level.

TCDB and Pfam use statistical and HMM sequence-based approaches, respectively, to establish homology. As previously discussed, direct applications of the TCDB and Pfam approaches to elevator transporters are not capable of producing meaningful assessments of homology due to the topological differences present in elevator transport domains. These approaches also require analysis of hundreds of sequences to robustly detect distant homology. In future work, we plan to build the complex computational machinery required to meaningfully reapply the TCDB and Pfam approaches to the hundreds of known elevator transporter sequences. For now, however, we restrict our sequence-based analysis to the limited set of of elevator transporters with at least one structure deposited in the PDB.

In the simplest model of sequence divergence over time, mutation occurs at a constant rate over time, all mutations (from any one amino acid to any other) are equally likely (for the sake of simplicity), and all mutations are independent. With these conditions, the probability P that a residue within a sequence has an amino acid identity different from its original identity by time t is given by

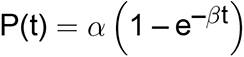

where *α* is a constant representing the probability that a mutation is not a back mutation, and *β* is a constant representing the mutation rate. This formula can be derived by extending an early model^24, 25^ of sequence divergence to account for back mutations or by generalizing the Jukes-Cantor model^26^ of nucleic acid sequence divergence to amino acids.

Since architecture is conserved far longer during protein evolution than sequence, we suggest that structural divergence (as measured by TM-score) can stand in for time in the equation above, at least to a first approximation. With this premise, we have found that the above simple model of evolution effectively captures the relationship between sequence identity and structural similarity in elevator transport domains (Fig. 4C). The existence of this relationship shows that the amount of sequence divergence present in elevator transport domains is consistent with the idea that these elevator transporters evolved from a common ancestor. We also note that this same relationship between sequence identity and structural similarity is present among homologous proteins in CATH families.^27^

#### 2.4.3 Structural Complexity

Structural complexity is difficult to quantify in absolute terms, but we suggest that the elevator transport domains analyzed here are structurally complex in qualitative terms, specifically relative to protein motifs in the MALISAM^28^ database. MALISAM is the foremost database of protein motifs that have been identified as analogous, meaning the pairs of proteins share striking structural similarities but have been proven to share no common ancestor. The analogous motifs in the MALISAM database consist of up to 100 residues (including loops) folded into 4 to 6 *α*-helices or *β*-strands arranged into simple planar or box-shaped architectures (Fig. 4E). By comparison, the transport domains consist of about 150 residues (excluding loops) folded into 10 *α*-helices arranged in a considerably more complex overall architecture.

The MALISAM database provides the best insight available into the degree of complexity generally present in analogous protein structures formed through convergent evolution. The substantially higher relative complexity of the transport domains is consistent with the idea that they formed through divergent evolution. Taken together with our analysis of the degrees of structural, topology-corrected sequence, and functional similarity present among the transport domains, we suggest that these results form a compelling body of evidence that indicates elevator transporters are homologous.

### 2.5 Structure-Based Phylogenetic Tree

Using our pairwise TM-score measurements, we constructed a phylogenetic tree (Fig. 5). In the tree, transporters from the same TCDB family group together into their own clades, and the tree further groups TCDB families from the same superfamily into larger clades. This result makes sense, as our analysis has revealed that transporters in the same TCDB family share common structural features, namely a particular transport domain topology and oligomerization state (Table 1). Overall, the tree is consistent with the TCDB classification system but also offers a higher level organization for it.

**Figure 5:**
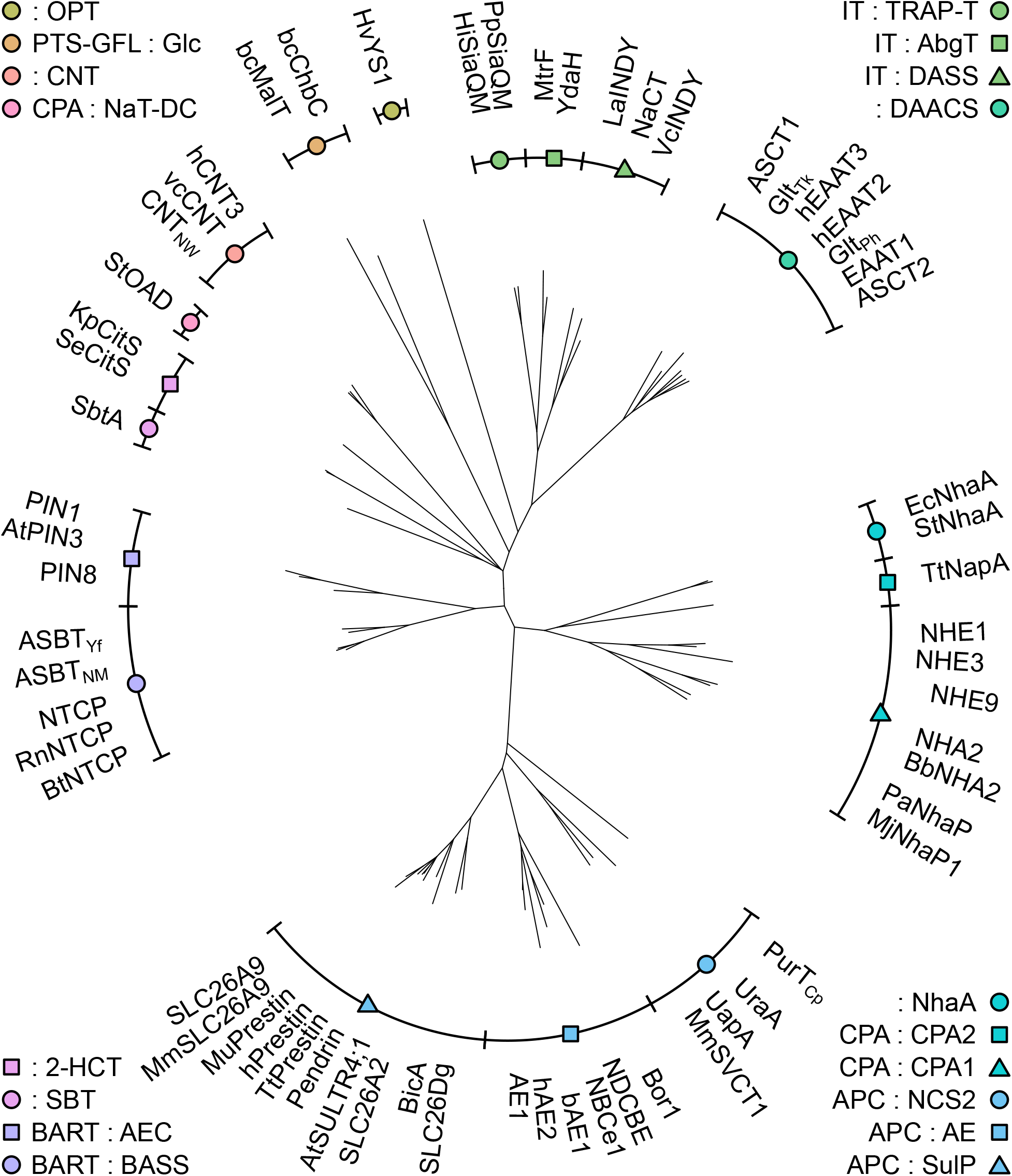
Phylogenetic tree of elevator transporters. The tree was produced by applying neighbor joining to the distance matrix generated from pairwise structural comparisons of the elevator transport domains. The tree separates TCDB families and superfamilies into their own clades, making the classification consistent with TCDB while offering a higher level organization.

In practice, the phylogenetic tree enables visualization and quantitative measurement of structural and evolutionary relationships among elevator transporters and their families. It offers a natural framework for new investigations into the structural and functional features that can be shared among elevator transporters and into how broadly these features are shared. By extension, the tree is also a natural tool for suggesting possibilities for yet to be characterized structural and functional features in elevator transporters. Indeed, in preliminary work, we have already found several examples of features that may be shared among elevator transporters, including domain movements, helix flexibility, and ion binding sites.

#### 2.5.1 Comparing Domain Movements

In elevator transporters, transport is primarily achieved through rigid-body movement of the transport domain relative to the scaffold domain. In previous work,^29^ we developed system-specific collective variables^30^ (i.e., formal mathematical functions) describing the domain movements involved in the transport mechanism of the elevator transporter LaINDY. Using our annotation of the transport domain helices of elevator transporters, we can directly generalize our original collective variable definitions for use with any elevator transporter, offering a new framework for rigorous quantitative comparisons of transport domain movements.

In describing the elevator mechanism of LaINDY, we defined two collective variables: z for the translation and *θ* for the rotation of the transport domain relative to the scaffold domain (Fig. 6A), and we calculated that the LaINDY OF structure (PDB `6WTW`) resides at z = 4.2 Å, *θ* = 19.5*^◦^*. Exploiting the internal structural symmetry (i.e., inverted repeats) in LaINDY and in the definitions of the collective variables z and *θ*, we predicted that the IF state of LaINDY would reside at z = –4.2 Å, *θ* = –19.5*^◦^*.^29^ Using the same arguments and protocols, we have defined equivalent collective variables for the other structurally characterized transporters in the IT : DASS family, namely VcINDY and NaCT. We have found that the VcINDY IF structure (PDB `5UL7`) resides at z = –8.3 Å, *θ* = –22.1*^◦^*, and the NaCT IF structure (PDB `7JSK`) resides at z = –6.3 Å, *θ* = –14.1*^◦^*.

**Figure 6:**
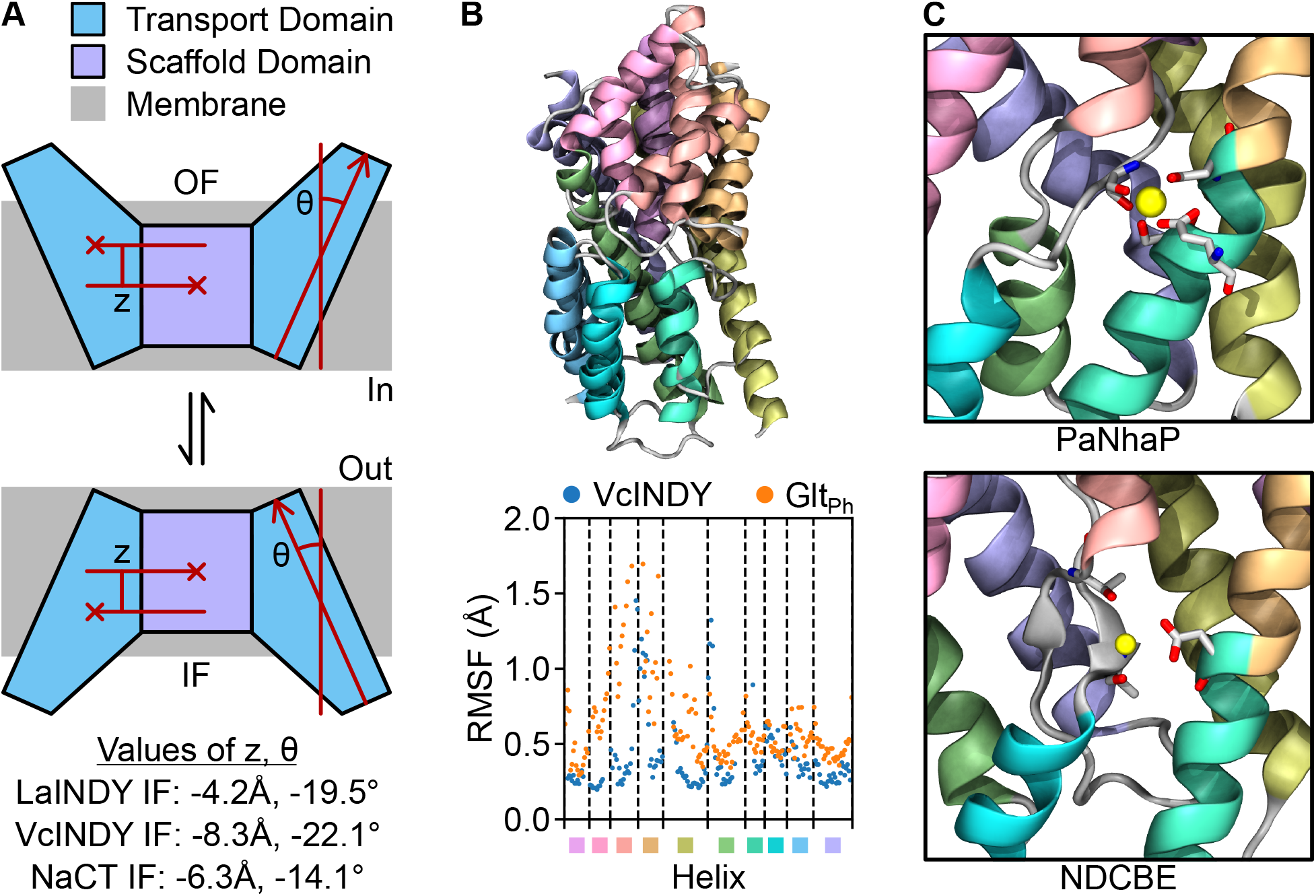
Examples of structural and functional features that may be shared among elevator transporters: domain movements, helix flexibility, and ion binding sites. **(A)** In the elevator transport mechanism, the IF *^__^s;-* OF transition involves translation and rotation of the transport domain relative to the scaffold domain, depicted schematically as changes in z and *θ*. In past work, we have developed rigorous mathematical definitions for z and *θ* which can be directly generalized to any elevator transporter. Analysis of LaINDY, VcINDY, and NaCT illustrates the range of z and *θ* values associated with the IF state in the IT : DASS family. We speculate that the z and *θ* values associated with equivalent functional states will generally be more similar for transporters within the same clade or in adjacent clades in the phylogenetic tree than they will be for more distantly related transporters. **(B)** Comparison of the transport domain helix flexibility in VcINDY and Glt_Ph_. (Top) Superposition of the transport domains of VcINDY (PDB `5UL9-A`) and Glt_Ph_ (PDB `7RCP-A`). (Bottom) Plot of the root-mean-square fluctuation (RMSF) of structurally equivalent transport domain helix C*_α_* atoms from VcINDY and Glt_Ph_. RMSF values were calculated for each transporter after superposing the ensemble of all transport domain structures deposited in the PDB (44 protomers from 13 PDBs for VcINDY and 122 protomers from 44 PDBs for Glt_Ph_). For helices of VcINDY and Glt_Ph_ that lie on top of one another when structurally superposed, there is generally good agreement in the corresponding RMSF values. For helices that are less structurally similar, specifically the upper left (pink) and upper center (red) helices, the corresponding RMSF values are also less similar. **(C)** Comparison of ion binding sites from elevator transporters PaNhaP (PDB `4CZA-B`) and NDCBE (PDB `7RTM-B`). The transport domain topologies of these two transporters are different, but the two ion binding sites are formed by residues from equivalent secondary structural elements. We speculate that the conserved architecture of the transport domains enables convergent evolution to form structurally similar ion binding sites, suggesting that known elevator transporter ion binding sites can be used to propose possible locations for unresolved ion binding sites in other elevator transporters.

Given the necessarily deep connection between structure and the extent of possible domain movement, we speculate that closely related (i.e., structurally similar) elevator transporters will undergo similar domain movements during transport. Our analysis of the IT : DASS family transporters gives one example of the range of transport domain positions associated with a particular functional state within a family. As we perform more comprehensive analysis of more distantly related elevator transporters, it will be interesting to see what patterns emerge and how they correspond to the phylogenetic tree.

#### 2.5.2 Investigating Conserved Helix Flexibility

While elevator transport is primarily achieved through rigid-body domain movements, small helix movements within the transport domain are also involved in substrate and ion binding in elevator transporters. For example, the upper center (red) and upper right (orange) transport domain helices form a hairpin in Glt_Ph_, which has been observed, both experimentally and computationally, to become more flexible when the substrate unbinds.^31, 32^ These same helices also form a hairpin in VcINDY (which belongs to a clade adjacent to Glt_Ph_ in the phylogenetic tree), and recently published structures have shown that this hairpin also becomes more flexible when Na^+^ unbinds.^33^ Interestingly, superposition of Glt_Ph_ and VcINDY reveals that the local packing of these hairpin helices is different for these two transporters, and we speculate that these local structural differences result in significant differences in the relative flexibility of these two hairpins (Fig. 6B). Overall, we suggest that similar functionally relevant helix movements may be conserved among related elevator transporters, and the phylogenetic tree offers a guide for investigating this conservation.

#### 2.5.3 Identifying Related Ion Binding Sites

Finally, in further preliminary analysis, we have also observed structurally similar ion binding sites in PaNhaP and NDCBE (Fig. 6C). These transporters are from different TCDB families and have different transport domain topologies. Nevertheless, the two binding sites are formed by residues from equivalent secondary structural elements in the conserved architecture shared by both transport domains. We speculate that these binding sites were not inherited from a common ancestor but rather developed independently through convergent evolution operating on the conserved transport domain architecture. Regardless of the evolutionary mechanism by which these ion binding sites were formed, this result suggests that distantly related transporters may be used to suggest possibilities for the locations of unknown ion or substrate binding sites.

### 2.6 Scaffold Domain

During the IF *^__^s;-* OF transition, the transport domain of elevator transporters undergoes a large-scale, rigid-body motion relative to the scaffold domain. A structure may capture a transporter at any point within this transition, so it was necessary to treat the transport and scaffold domains separately to perform meaningful structural superpositions and comparisons with TM-align. Individual domains within a protein are also known to experience individual evolutionary pressures,^7, 19–21^ and this information also justified treating the domains separately. We chose to focus our analysis first and primarily on the transport domains because the structural similarity among them was striking and immediately apparent. However, previous work has noted that there are structurally similar features in the scaffold domains of elevator transporters as well.^3, 11^ Through comprehensive analysis of the available elevator transporter structures, we have also observed a highly conserved architecture and topology in the scaffold domain (Fig. 7).

**Figure 7:**
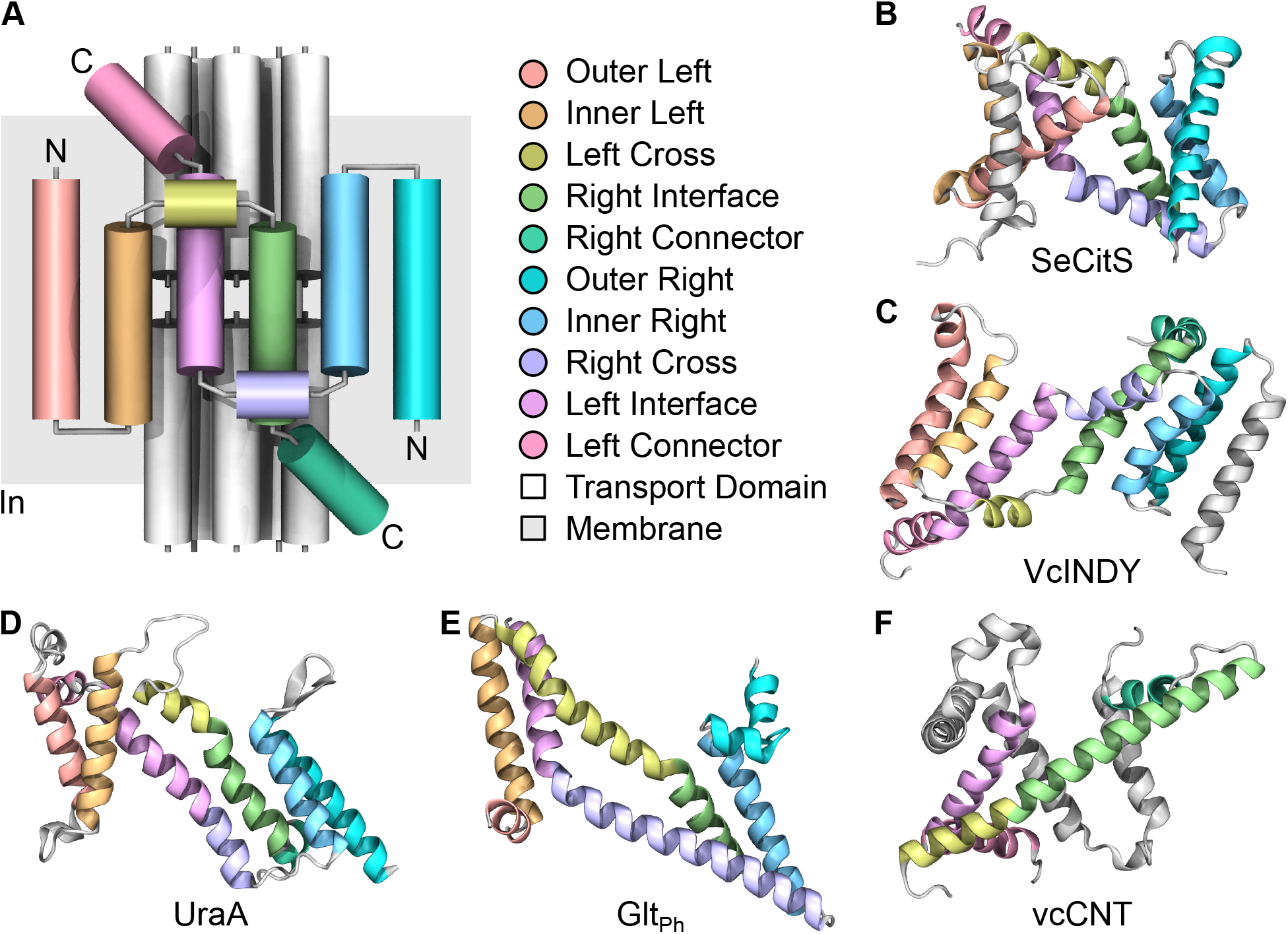
The architecture and topology of the scaffold domain of elevator transporters. **(A)** A simplified schematic of the scaffold domain, with each of the 10 constituent helices assigned a unique name and color. **(B)** The scaffold domain of SeCitS (PDB `5A1S-A`), which demonstrates close adherence to the idealized scaffold domain template from (A). **(C)** The scaffold domain of VcINDY (PDB `5UL9-A`), which also demonstrates close adherence to the template, except its chirality is reversed. All members of the IT superfamily exhibit this reversed chirality. **(D)** The scaffold domain of UraA (PDB `5XLS-A`). The N-to-C direction and ordering of the helices is reversed relative to the template, but the template architecture is maintained nevertheless. All members of the APC superfamily exhibit this reversed N-to-C helix direction and ordering. **(E)** The scaffold domain of Glt_Ph_ (PDB `7RCP-A`). This scaffold domain exhibits increased architectural variability relative to the others, though its correspondence with the template is still apparent. **(F)** The scaffold domain of vcCNT (PDB `4PD6-A`). This scaffold domain exhibits even greater architectural variability relative to the others, and its correspondence with the template is difficult to discern, except through direct comparison with the scaffold domain of VcINDY and other members of the IT superfamily. Members of the CNT family (like vcCNT) and the PTS-GFL : Glc family exhibit this level of architectural variability in their scaffold domains.

Within the conserved architecture and topology of the scaffold domain, two helices at the center of the domain form the primary interface with the transport domain. These interfacial helices are each directly attached to a “connector” helix that attach to the transport domain. The other ends of the interfacial helices connect to the rest of the scaffold domain helices in a complex intertwined pattern. Overall, the conserved scaffold domain topology also contains an inverted repeat pattern.

Despite there being enough structural similarity in the scaffold domains to develop this structural template, the overall structural variability in the scaffold domains is far greater than in the transport domains. The outer helices (in our template nomenclature) are often absent or joined to extra helices not present in our template, and they rearrange themselves significantly in different transporter families, enabling different oligomerization states. More significantly, the chirality of the IT superfamily scaffold domain is opposite that of the other scaffold domains, and the N-to-C direction of the APC superfamily scaffold domain helices is reversed relative to the other scaffold domains. Further, the scaffold domains of the CNT and PTS-GFL : Glc families are so different from the template that we would not recognize them as following that template were it not for direct comparison with the scaffold domains of closely related transporter families.

Because of the overall structural variability in the scaffold domains, we do not believe structural superposition and meaningful quantitative comparison of these domains are possible with TM-align. Still, noting that structural similarity is present throughout the entirety of elevator transporters is important. The search for shared functional structural features in distantly related elevator transporters need not be limited to the transport domain alone, and this observation also substantially increases the overall complexity of the structural similarity, strengthening the case that these proteins are homologous. This observation also explains how Pfam has been able to form the CPA/AT clan, which has detected at the sequence level a homologous relationship among several elevator transporter families with different transport domain topologies (specifically the CPA : NaT-DC, 2-HCT, SBT, BART : AEC, BART : BASS, NhaA, and CPA : CPA1 families).

We also suggest that the structural variability of the scaffold domains is already represented, at least in part, in the phylogenetic tree derived from the transport domains. Despite containing no information from the scaffold domains, the phylogenetic tree separates elevator transporters with different oligomeric states into distinct clades, even when the transport domain topology of the transporters is the same. This makes complete sense in light of the close structural and functional coupling between the transport and scaffold domains. Structural variability in the scaffold domain generally must generate a structural response in the transport domain and vice versa for the transporter to remain operable.

## 3 Conclusion

Through comprehensive examination of the pertinent structures available in the PDB, we have found that 62 elevator transporters from 18 TCDB families share a conserved architecture in their transport domains, which consists of 10 helices connected to each other in 8 distinct topologies. They also share a conserved architecture in their scaffold domains, though there is a higher degree of structural variability than in the transport domains. Through detailed, topology-corrected analysis of the similarity in structure, sequence, and function of the transport domains, we have found compelling evidence that the elevator transporters are homologs that evolved from a common ancestor. With this context, we constructed a phylogenetic tree for the elevator transporters which allowed us to visualize and quantify the evolutionary and structural relationships among them and their families. The tree is useful for guiding new investigation into how broadly functional structural features are shared, and we have already found examples of similar shared domain and helix motions and ion binding sites in preliminary work.

In this study, we used structural similarity to infer phylogeny, and this approach necessarily assumes that there is a certain amount of structural rigidity in the transport domains of elevator transporters. Within the elevator transport mechanism, the transport domain generally moves as a rigid body relative to the scaffold domain, but small helix rearrangements within the transport domain have been noted during substrate and ion binding for, e.g., Glt_Ph_^31, 32^ and VcINDY.^33^ To test the validity of the assumption of rigidity, we superposed all of the available transport domain structures individually for each elevator transporter, and we observed only small helix rearrangements relative to the larger rearrangements observed when comparing different transport domains (Fig. 2B). This result validates the assumption of structural rigidity in our approach; however, in future work, the estimates of evolutionary distance between proteins in the phylogenetic tree could be improved by using an ensemble of structures derived from experiments and/or molecular dynamics simulations of each protein.

Like the elevator mechanism, the rocking bundle mechanism is one of the handful of canonical mechanisms utilized to achieve alternating access, and it is used by a wide variety of biomedically significant transporters.^3^ In this mechanism, a core domain rocks back and forth against a stationary scaffold domain, alternatively exposing the central substrate binding pocket to one side of the membrane and then the other.^34, 35^ Within the TCDB system, the APC superfamily contains all known families of rocking bundle transporters as well as some families of elevator transporters,^36, 37^ indicating that there is strong sequence-based evidence of homology between elevator and rocking bundle transporters. Pioneering work has also noted structural similarity in the scaffold domains of elevator and rocking bundle transporters,^3, 11^ and we note that this similarity extends to the transport/core domains as well. In future work, we plan to apply our approach to the rocking bundle transporters, creating a unified phylogenetic tree for all rocking bundle and elevator transporters.

## 4 Methods

### 4.1 Identification of All Available Elevator Transporter Structures

To identify the structures of all elevator transporters that have been deposited in the PDB, we started with the collection of elevator transporter structures identified in a recent review,^5^ and we augmented it with additional elevator transporter structures identified through our own literature review. In compiling our collection of elevator transporter structures, we did not include structures of any energy coupling factor (ECF)-type adenosine triphosphate (ATP)-binding cassette (ABC) transporters, nor did we include structures of the transporters TmPiT (PDB `6L85`), BbZIP (PDB `5TSA`), or TtCcdA (PDB `5VKV`). These transporters are sometimes classified as elevator transporters, but we consider their modes of transport to be fundamentally distinct from the canonical elevator transport mechanism established by comparison of the first IF and OF structures of Glt_Ph_.^4^ Finally, we also excluded the structure of the transporter Bor1p (PDB `5SV9`) due to its very low resolution (5.90 Å).

From this process, we identified a collection of structures for 68 elevator transporters that have been released in the PDB as of May 10, 2023. As a representative structure of each transporter, we selected for additional analysis the structure with the highest overall resolution. Within each of these structures, we further selected the protomer assigned the alphabetically first protein chain (usually chain A).

In the elevator transport mechanism, the transport domain moves significantly with respect to the scaffold domain during the IF *^__^s;-* OF transition, and individual solved structures may capture a transporter at any point within this transition. As such, to quantitatively assess structural similarity among elevator transporters, we manually decomposed each transporter into its transport and scaffold domains and considered them separately, focusing in this work primarily on the transport domains.

### 4.2 Annotation of Transport Domain Helices

Within the structures of the transport domains, we noticed a widely conserved architecture that consisted of 10 helices (Fig. 2). We found that only 6 of the transport domains did not contain this conserved architecture, namely those from ecUlaA (PDB `4RP9`), pmU-laA (PDB `5ZOV`), ecManYZ (PDB `7DYR`), lmManYZ (PDB `7VLY`), lsManYZ (PDB `7XNO`), and llManYZ (PDB `8HFS`), and we excluded them from further analysis. This left us with a set of 62 elevator transporters from 18 TCDB families (Table 1). For each of these elevator transporters, we then manually annotated its transport domain helices by compiling a list of the residues that constituted each helix in the conserved architecture (Table S1). We developed this list using secondary structure assignments produced by STRIDE^38^ augmented by visual inspection in VMD.^39^

Most elevator transport domains are composed entirely of one set of inverted repeats, resulting in complete structural symmetry between the upper and lower halves of the transport domains (Fig. 3). This symmetry complicated helix annotation as two structural superpositions were possible for each pair of transport domains. Ultimately, we decided to use the orientations of the transport domains relative to the membrane to make the superpositions functionally consistent.

### 4.3 Structural Comparison of Transport Domains

We next sought to quantify the structural similarity in the transport domains using TM-align.^17^ The basic function of TM-align is to automatically identify the optimal superposition of two structures such that a quantitative measure of structural similarity called TM-score^18^ is maximized. To find this optimal superposition and calculate the TM-score, TM-align identifies a one-to-one mapping of atoms from the two structures that occupy the same region in space after being structurally superposed. An inherent limitation of the TM-align algorithm is that structurally equivalent atoms must appear in the same order within the two polypeptide sequences to be included in the TM-align mapping.

Within the elevator transport domains, the architecture was widely conserved, but the topology was not (Fig. 3). Specifically, the 10 helices of the conserved architecture were ordered in different ways within the polypeptide sequence, and the N-to-C orientations relative to the membrane normal axis were different for equivalent helices. Thus, to use TM-align to perform global structural superposition and comparison of the transport domains, we needed to first correct for these topological differences, which we did by using VMD to produce a modified PDB file for each transport domain.

To generate each modified PDB file, we selected the C*_α_* atoms of each residue previously annotated as belonging to the 10 conserved helices of the shared transport domain architecture. We then renumbered the residues such that the conserved helices appeared in a consistent order in the modified PDB files and such that the N-to-C orientations of the conserved helices were consistent relative to the membrane normal axis. We finally output a modified PDB file for each transport domain consisting only of helix C*_α_* atoms with their modified residue ordering.

Using the modified PDB files, we then performed pairwise structural comparisons of the transport domains with TM-align. With the reordered residues from the modified PDB files, TM-align was able to generate a global mapping of structurally equivalent residues for each pair of transport domain structures. From this global mapping, TM-align automatically quantified the pairwise structural similarity between the transport domains via both the TM-score and the root-mean-square displacement (RMSD) of the equivalent C*_α_* atoms, and it also produced a structure-based sequence alignment for each pair of structures.

### 4.4 Construction of the Phylogenetic Tree

To visualize the evolutionary relationships among the elevator transport domains, we next generated a phylogenetic tree. To do this, we first converted our previously calculated pairwise TM-score measurements to a distance matrix by subtracted them from 1, which was appropriate because TM-score is a measure of similarity that ranges from 0 to 1. We then applied the classic neighbor joining algorithm^40^ to the distance matrix using the T-REX web server^41^ to generate the phylogenetic tree, and we visualized the resulting tree using iTOL.^42^

## 5 Data Availability

The principal elevator transporter structures used in this study are available from the PDB under accession codes `7RCP` (Glt_Ph_), `5DWY` (Glt_Tk_), `5LM4` (EAAT1), `7XR4` (hEAAT2), `8CUA` (hEAAT3), `7P4I` (ASCT1), `7BCT` (ASCT2), `5UL9` (VcINDY), `6WTW` (LaINDY), `7JSK` (NaCT), `4R1I` (MtrF), `4R0C` (YdaH), `7QE5` (HiSiaQM), `7QHA` (PpSiaQM), `7WST` (HvYS1), `3QNQ` (bcChbC), `5IWS` (bcMalT), `4PD6` (vcCNT), `5L26` (CNT_NW_), `6KSW` (hCNT3), `6IWW` (StOAD), `5XAS` (KpCitS), `5A1S` (SeCitS), `7EGK` (SbtA), `7Y9T` (PIN1), `7WKW` (AtPIN3), `7QP9` (PIN8), `3ZUX` (ASBT_NM_), `6LGV` (ASBT_Yf_), `7VAF` (RnNTCP), `7VAE` (BtNTCP), `7ZYI` (NTCP), `7S24` (EcNhaA), `7A0W` (StNhaA), `5BZ3` (TtNapA), `4CZB` (MjNhaP1), `4CZ8` (PaNhaP), `7DSW` (NHE1), `7X2U` (NHE3), `6Z3Z` (NHE9), `7P1J` (BbNHA2), `7B4L` (NHA2), `5XLS` (UraA), `7TAK` (PurT_Cp_), `5I6C` (UapA), `7YTW` (MmSVCT1), `5L25` (Bor1), `7RTM` (NDCBE), `8D9N` (bAE1), `7UZ3` (AE1), `8GVC` (hAE2), `6CAA` (NBCe1), `6KI1` (BicA), `7LHV` (AtSULTR4;1), `5DA0` (SLC26Dg), `7XLM` (SLC26A2), `6RTC` (MmSLC26A9), `7CH1` (SLC26A9), `7SUN` (MuPrestin), `7S8X` (TtPrestin), `7LGU` (hPrestin), and `7WK1` (Pendrin).

Additionally, the VcINDY structure used in Fig. 1A is available from the PDB under accession code `5UL7`. The structures used to generate the Glt_Ph_ superposition in Fig. 2B and Fig. 6B are available from the PDB under accession codes `1XFH`, `2NWL`, `2NWW`, `2NWX`, `3KBC`, `3V8F`, `3V8G`, `4IZM`, `4OYE`, `4OYF`, `4P19`, `4P1A`, `4P3J`, `4P6H`, `4X2S`, `5CFY`, `6BAT`, `6BAU`, `6BAV`, `6BMI`, `6CTF`, `6UWF`, `6UWL`, `6V8G`, `6WYJ`, `6WYK`, `6WYL`, `6WZB`, `6X01`, `6X12`, `6X13`, `6X14`, `6X15`, `6X16`, `6X17`, `7AHK`, `7RCP`, `7UG0`, `7UGD`, `7UGJ`, `7UGV`, `7UGX`, `7UH3`, and `7UH6`. The aligned structural analogs used in Fig. 4E are available from the MALISAM database under entry `d1b05a_d1nkia_`. The structures used to generate the VcINDY superposition in Fig. 6B are available from the PDB under accession codes `4F35`, `5UL7`, `5UL9`, `5ULD`, `5ULE`, `6OKZ`, `6OL0`, `6OL1`, `6WTX`, `6WU3`, `6WW5`, `7T9F`, and `7T9G`. The PaNhaP structure used in Fig. 6C is available from the PDB under accession code `4CZA`.

Finally, the topology-corrected PDB files that were compared pairwise by TM-align, the mappings from the residue numbers in the topology-corrected PDB files to the residue numbers in the original PDB files, the log files that were generated during the pairwise comparisons by TM-align, and PDB files aligned to comprehensively demonstrate the pairwise structural superpositions produced by TM-align are all available on Zenodo (doi.org/10.5281/zenodo.8040643). All other data generated or analyzed in this study are included in this published article and its supplementary information files.

## 6 Acknowledgements

Research reported in this publication was supported by grants from the National Institutes of Health under award numbers P41-GM104601, R24-GM145965, and R01-DK135088. NT acknowledges support from the National Science Foundation Graduate Research Fellowship Program under Grant No. 1746047. Any opinions, findings, and conclusions or recommendations expressed in this material are those of the authors and do not necessarily reflect the views of the National Institutes of Health or the National Science Foundation. The authors gratefully acknowledge helpful discussions with Olga Boudker.

## 7 Author Contributions

NT conceived, designed, and executed the study, and ET supervised the work. NT prepared the initial draft of the manuscript, and ET and NT revised it together.

