## Supplementary material for "Structure Reveals Homology in Elevator Transporters": Table S1

| Protein | Upper Core (Mauve) | Upper Left (Pink) | Upper Center (Red) | Upper Right (Orange) | Right Backbone (Yellow) | Lower Core (Green) | Lower Right (Turquoise) | Lower Center (Cyan) | Lower Left (Blue) | Left Backbone (Violet) |
|---|---|---|---|---|---|---|---|---|---|---|
| $Glt_{Ph}$ | 312-329 | 377-389 | 358-370 | 335-351 | 75-106 | 390-416 | 296-309 | 279-292 | 258-275 | 227-254 |
| $Glt_{Tk}$ | 315-332 | 382-393 | 361-373 | 338-352 | 77-109 | 394-420 | 299-313 | 280-295 | 260-277 | 228-256 |
| EAAT1 | 380-395 | 442-451 | 426-438 | 402-419 | 113-143 | 452-479 | 364-377 | 346-358 | 325-342 | 295-321 |
| hEAAT2 | 399-416 | 461-470 | 445-457 | 422-438 | 115-142 | 471-496 | 383-396 | 365-379 | 344-361 | 306-340 |
| hEAAT3 | 368-385 | 432-439 | 414-426 | 391-404 | 86-116 | 440-465 | 352-365 | 334-348 | 313-330 | 276-308 |
| ASCT1 | 380-397 | 443-451 | 426-438 | 403-416 | 113-139 | 452-480 | 366-377 | 345-359 | 325-342 | 287-321 |
| ASCT2 | 388-403 | 452-459 | 437-446 | 411-428 | 121-153 | 460-488 | 372-385 | 353-367 | 333-350 | 303-329 |
| VcINDY | 400-412 | 201-210 | 378-396 | 358-375 | 435-459 | 175-193 | 422-429 | 151-166 | 131-147 | 214-238 |
| LaINDY | 420-434 | 211-223 | 399-416 | 377-394 | 456-485 | 188-205 | 443-449 | 156-175 | 135-150 | 230-255 |
| NaCT | 487-499 | 228-240 | 465-483 | 445-460 | 522-552 | 202-220 | 509-516 | 141-158 | 121-137 | 250-276 |
| MtrF | 438-452 | 191-205 | 417-434 | 397-411 | 480-505 | 168-181 | 461-470 | 147-164 | 128-142 | 218-241 |
| YdaH | 418-432 | 178-194 | 396-414 | 375-392 | 458-486 | 155-169 | 440-452 | 134-151 | 115-131 | 205-233 |
| HiSiaQM | 544-560 | 344-353 | 522-540 | 502-518 | 579-602 | 323-339 | 566-575 | 301-319 | 282-298 | 357-386 |
| PpSiaQM | 352-367 | 152-161 | 330-346 | 310-325 | 388-410 | 131-148 | 374-383 | 109-126 | 91-103 | 165-194 |
| HvYS1 | 470-501 | 546-559 | 449-464 | 421-445 | 165-192 | 112-149 | 199-212 | 81-103 | 51-73 | 505-533 |
| bcChbC | 286-293 | 386-394 | 251-275 | 231-247 | 346-369 | 300-311 | 333-340 | 315-330 | Absent | 398-431 |
| bcMalT | 308-315 | 395-407 | 240-249, 271-288 | 221-237 | 366-382 | 317-328 | 354-359 | 333-351 | Absent | 410-439 |
| vcCNT | 353-363 | 186-195 | 332-343 | 316-329 | 386-415 | 170-182 | 369-382 | 155-168 | 140-150 | 199-220 |
| $CNT_{NW}$ | 353-363 | 186-195 | 332-342 | 314-330 | 386-418 | 170-182 | 369-382 | 155-168 | 140-150 | 199-220 |
| hCNT3 | 551-560 | 375-383 | 520-535 | 504-517 | 585-613 | 358-370 | 568-579 | 343-356 | 328-338 | 387-408 |
| StOAD | 190-201 | 218-232 | 204-214 | 420-431 | 158-183 | 374-379 | 408-419 | 385-398 | 233-244 | 337-364 |
| KpCitS | 170-182 | 205-222 | 186-201 | 435-446 | 145-166 | 388-399 | 417-434 | 403-414 | 223-236 | 360-384 |
| SeCitS | 170-182 | 205-222 | 186-201 | 435-447 | 145-166 | 388-400 | 417-434 | 403-414 | 223-236 | 360-384 |
| SbtA | 310-321 | 136-146 | 115-128 | 357-370 | 278-304 | 100-111 | 341-356 | 327-335 | 147-164 | 71-92 |
| PIN1 | 567-577 | 128-143 | 114-125 | 610-621 | 537-562 | 98-110 | 598-609 | 582-591 | 146-164 | 70-92 |
| AtPIN3 | 584-595 | 127-143 | 114-125 | 628-640 | 555-580 | 98-109 | 613-627 | 600-609 | 144-164 | 69-92 |
| PIN8 | 311-321 | 132-148 | 119-130 | 355-366 | 282-307 | 102-114 | 340-354 | 327-336 | 149-163 | 69-92 |
| $ASBT_{NM}$ | 96-107 | 281-297 | 266-277 | 140-151 | 67-92 | 251-262 | 125-139 | 112-121 | 298-308 | 222-247 |
| $ASBT_{Yf}$ | 90-101 | 273-289 | 260-271 | 134-144 | 61-86 | 245-256 | 119-133 | 108-115 | 290-299 | 216-241 |
| RnNTCP | 87-98 | 280-296 | 263-273 | 131-142 | 58-83 | 248-259 | 116-130 | 103-112 | 297-310 | 219-244 |
| BtNTCP | 87-98 | 280-296 | 263-273 | 131-142 | 58-83 | 248-259 | 116-130 | 103-112 | 297-310 | 219-244 |
| NTCP | 87-98 | 280-296 | 263-273 | 131-142 | 58-82 | 248-259 | 116-130 | 103-112 | 297-310 | 219-243 |
| EcNhaA | 121-131 | 355-372 | 338-351 | 165-175 | 91-117 | 325-336 | 150-164 | 134-144 | 373-384 | 287-314 |
| StNhaA | 121-131 | 355-372 | 338-351 | 165-175 | 91-117 | 325-336 | 150-164 | 134-144 | 373-381 | 287-314 |

Table continued on next page.

Table continued from previous page.

| Protein | Upper Core (Mauve) | Upper Left (Pink) | Upper Center (Red) | Upper Right (Orange) | Right Backbone (Yellow) | Lower Core (Green) | Lower Right (Turquoise) | Lower Center (Cyan) | Lower Left (Blue) | Left Backbone (Violet) |
|---|---|---|---|---|---|---|---|---|---|---|
| TtNapA | 113-122 | 350-367 | 333-345 | 158-173 | 84-109 | 318-328 | 144-157 | 128-138 | 368-385 | 288-316 |
| MjNhaP1 | 119-129 | 379-404 | 349-367 | 162-175 | 85-112 | 335-344 | 147-161 | 133-142 | 405-411 | 305-328 |
| PaNhaP | 116-126 | 387-411 | 364-380 | 160-173 | 86-112 | 350-359 | 145-159 | 134-141 | 412-421 | 322-346 |
| NHE1 | 224-235 | 476-498 | 461-468 | 268-281 | 186-214 | 446-455 | 253-267 | 240-245 | 499-504 | 411-437 |
| NHE3 | 178-188 | 434-455 | 417-425 | 222-237 | 140-168 | 403-412 | 207-221 | 193-201 | 456-464 | 368-395 |
| NHE9 | 201-212 | 458-479 | 444-450 | 245-256 | 159-188 | 432-438 | 230-244 | 217-226 | 480-487 | 396-422 |
| BbNHA2 | 230-240 | 480-499 | 461-475 | 276-293 | 200-222 | 446-457 | 266-275 | 245-258 | 500-515 | 417-438 |
| NHA2 | 231-241 | 481-500 | 461-478 | 277-294 | 201-227 | 447-456 | 267-276 | 246-259 | 501-516 | 418-440 |
| UraA | 87-111 | 29-39 | 75-85 | 225-240 | 263-280 | 303-318 | 241-253 | 289-298 | 14-28 | 43-61 |
| $PurT_{Cp}$ | 108-131 | 44-57 | 91-102 | 256-275 | 300-317 | 349-364 | 276-290 | 329-340 | 33-43 | 61-81 |
| UapA | 186-206 | 91-104 | 155-171 | 342-355 | 375-398 | 421-436 | 356-368 | 407-416 | 76-90 | 109-131 |
| MmSVCT1 | 152-178 | 63-76 | 120-129 | 325-340 | 360-380 | 404-418 | 341-353 | 390-398 | 51-62 | 83-105 |
| Bor1 | 120-140 | 52-62 | 97-112 | 292-306 | 338-349 | 466-484 | 307-320 | 359-366 | 33-51 | 68-81 |
| NDCBE | 559-581 | 490-504 | 539-552 | 780-799 | 821-838 | 880-891 | 800-809 | 847-856 | 477-489 | 510-526 |
| bAE1 | 504-524 | 439-448 | 484-499 | 679-694 | 720-737 | 776-794 | 695-705 | 746-747 | 421-438 | 455-471 |
| AE1 | 486-507 | 417-431 | 466-482 | 661-680 | 702-719 | 760-773 | 681-690 | 728-738 | 403-416 | 437-454 |
| hAE2 | 789-810 | 720-733 | 769-785 | 991-1010 | 1032-1048 | 1091-1102 | 1011-1019 | 1058-1068 | 706-719 | 740-757 |
| NBCe1 | 506-527 | 437-447 | 486-499 | 735-753 | 775-792 | 834-844 | 754-761 | 802-810 | 423-436 | 459-472 |
| BicA | 90-112 | 27-37 | 71-86 | 240-257 | 276-293 | 316-334 | 258-270 | 303-311 | 12-26 | 41-58 |
| AtSULTR4;1 | 172-195 | 110-120 | 153-166 | 327-346 | 365-382 | 405-422 | 347-359 | 392-400 | 91-109 | 124-141 |
| SLC26Dg | 94-114 | 36-46 | 79-90 | 220-240 | 259-275 | 299-317 | 241-252 | 285-294 | 21-35 | 50-67 |
| SLC26A2 | 214-239 | 123-133 | 166-184 | 377-395 | 416-430 | 455-471 | 396-408 | 440-448 | 108-122 | 138-152 |
| MmSLC26A9 | 162-191 | 86-95 | 129-142 | 334-346 | 366-382 | 405-421 | 347-359 | 391-400 | 70-85 | 101-117 |
| SLC26A9 | 162-191 | 86-96 | 129-142 | 330-346 | 365-381 | 405-419 | 347-359 | 391-400 | 70-85 | 100-117 |
| MuPrestin | 168-196 | 95-105 | 138-151 | 336-352 | 371-387 | 411-427 | 353-365 | 397-406 | 79-94 | 109-126 |
| TtPrestin | 168-196 | 95-105 | 138-151 | 335-352 | 371-388 | 411-427 | 353-364 | 397-406 | 76-94 | 109-126 |
| hPrestin | 169-196 | 95-105 | 138-151 | 336-352 | 371-388 | 411-427 | 353-365 | 397-406 | 76-94 | 109-126 |
| Pendrin | 180-206 | 99-109 | 142-155 | 346-362 | 381-397 | 421-437 | 363-375 | 407-416 | 80-98 | 113-130 |

**Table S1: Table of residues manually annotated as belonging to each transport domain helix of each elevator transporter.** See Fig. 2 for a schematic of the layout of the helices in the conserved architecture of the transport domains.
